# Transcriptional profiling of aging tissues from female and male African turquoise killifish

**DOI:** 10.1101/2023.06.20.545766

**Authors:** Alan Xu, Bryan B. Teefy, Ryan J. Lu, Séverine Nozownik, Alexandra M. Tyers, Dario R. Valenzano, Bérénice A. Benayoun

## Abstract

The African turquoise killifish is an emerging vertebrate model organism with great potential for aging research due to its naturally short lifespan. Thus far, turquoise killifish aging ‘omic’ studies using RNA-seq have examined a single organ, single sex and/or evaluated samples from non-reference strains. Here, we describe a resource dataset of ribosomal RNA depleted RNA-seq libraries generated from the brain, heart, muscle, and spleen from both sexes, as well as young and old animals, in the reference GRZ turquoise killifish strain. We provide basic quality control steps and demonstrate the utility of our dataset by performing differential gene expression and gene ontology analyses by age and sex. Importantly, we show that age has a greater impact than sex on transcriptional landscapes across probed tissues. Finally, we confirm transcription of transposable elements (TEs), which are highly abundant and increase in expression with age in brain tissue. This dataset will be a useful resource for exploring gene and TE expression as a function of both age and sex in a powerful naturally short-lived vertebrate model.

## Background and Summary

Aging is a complex breakdown in the processes that facilitate organismal homeostasis. Importantly, aging has been shown to broadly impact the landscape of genomic regulation across tissues, sexes, and species^1, 2^. This includes not only differences in canonical gene expression, but also in the expression of transposable elements (TEs)^2–8^. TEs are mobile repetitive genetic elements that are typically silenced in young tissues but become de-repressed with age. By examining how gene and TE expression changes with age, we can better understand the processes driving the aging process.

An important variable to consider when conducting any type of aging research are the myriad effects of biological sex^9^. For example, longevity is sex-dimorphic in humans in which females consistently outlive males^10^. The same trend is common across many animal species and appears to hold for most mammals^11, 12^. Sex also affects the risk of developing age-related diseases with men at higher risk of coronary artery disease and women at higher risk of Alzheimer’s disease^13–15^. Sex may also influence aging through differential activity of transposable elements TEs^16, 17^. Indeed, TE de-repression was shown to correlate with decreased lifespan in transgenic flies with different copy numbers of the TE-rich Y chromosome^18^.

An emerging powerful model to study aging in vertebrates is the African turquoise killifish *Nothobranchius furzeri*^19–26^. The turquoise killifish is the shortest-lived vertebrate that can be bred in captivity, with a naturally short lifespan of 4-6 months. Moreover, it is relatively inexpensive to maintain compared to other traditional vertebrate model organisms (*e.g*. mice). Accumulating studies are using RNA-seq in the turquoise killifish to understand the effects of aging and aging interventions on many different tissues^27–31^. However, most of these studies have either focused on a single tissue, a single sex, or used a non-reference strain of turquoise killifish (*e.g*. MZM-0410)^27–31^. In addition, these studies have also focused on genic transcription, leaving little known about how TE transcription is regulated with aging in this species.

Here, we generated ribosomal-RNA depleted bulk RNA-seq datasets from young (6- weeks-old) and old (16-weeks-old) male and female GRZ strain turquoise killifish brain, heart, muscle, and spleen (n= 4-5 per sex) (Figure 1A, Table 1). We found strong age effects and mild sex-dimorphism in all sampled tissues. We performed differential gene expression in each tissue to identify genes and TE regulated by age or by sex, and observed that age is a larger driver of gene expression differences than sex in these tissues and conditions. Furthermore, we showed that TEs are highly expressed across tissues, even in a healthy context, and upregulated with age in the brain. Lastly, as a proof-of-principle, we perform gene ontology (GO) analysis to demonstrate a common aging signature across multiple tissues in the turquoise killifish, characterized by increased immune/inflammatory gene expression, consistent with previous findings in other species.

**Figure 1.**
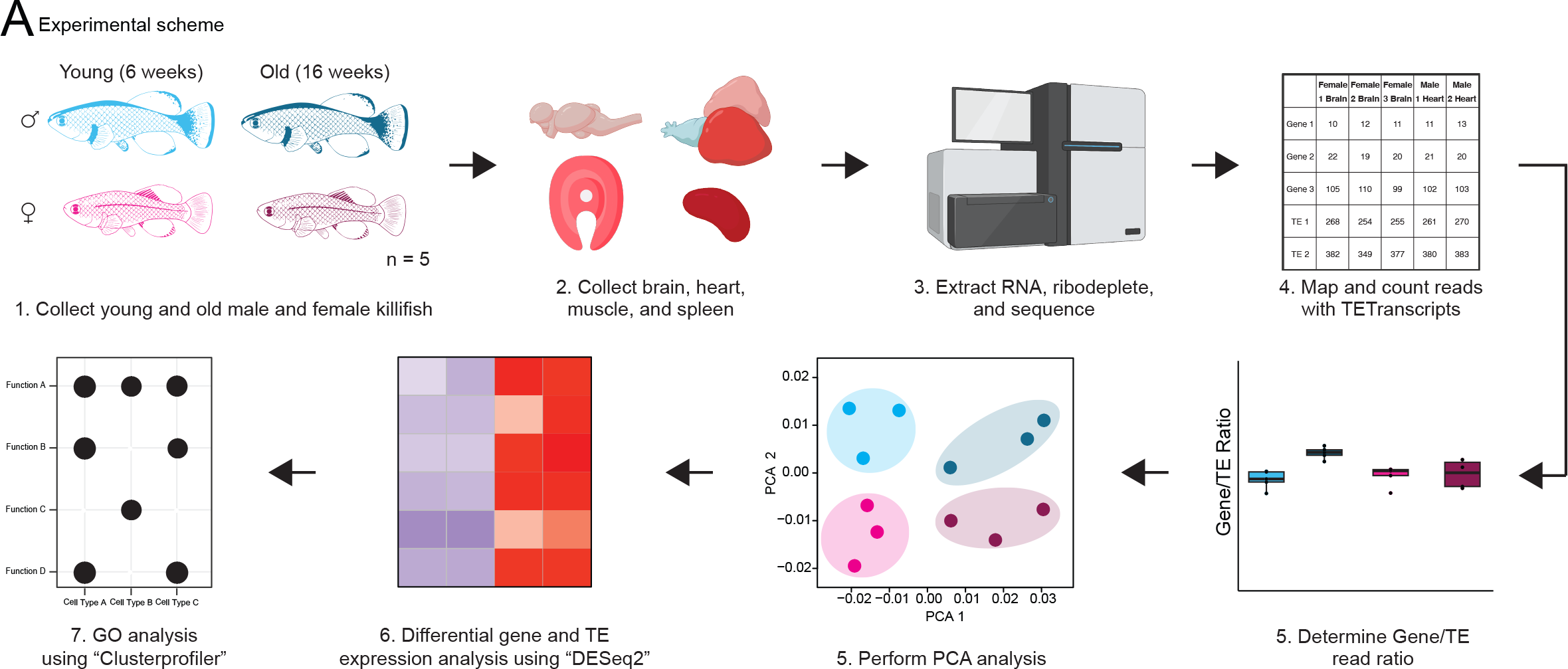
Experimental design and analytical pipeline. (A) The experimental design used to generate our RNA-seq dataset. Brain, heart, muscle, and spleen were dissected from sets of 5 young female, young male, old female, and old male GRZ strain killifish. RNA was extracted, depleted of ribosomal reads, and sequenced. After sequencing, reads were mapped to a turquoise killifish genome reference and counted with TETranscripts. The ratio of TE to gene reads was compared for each library, groups were contrasted for similarity using PCA analysis, differential gene expression was run using DESeq2, and gene ontology analysis was run using clusterProfiler.

**Table 1.**
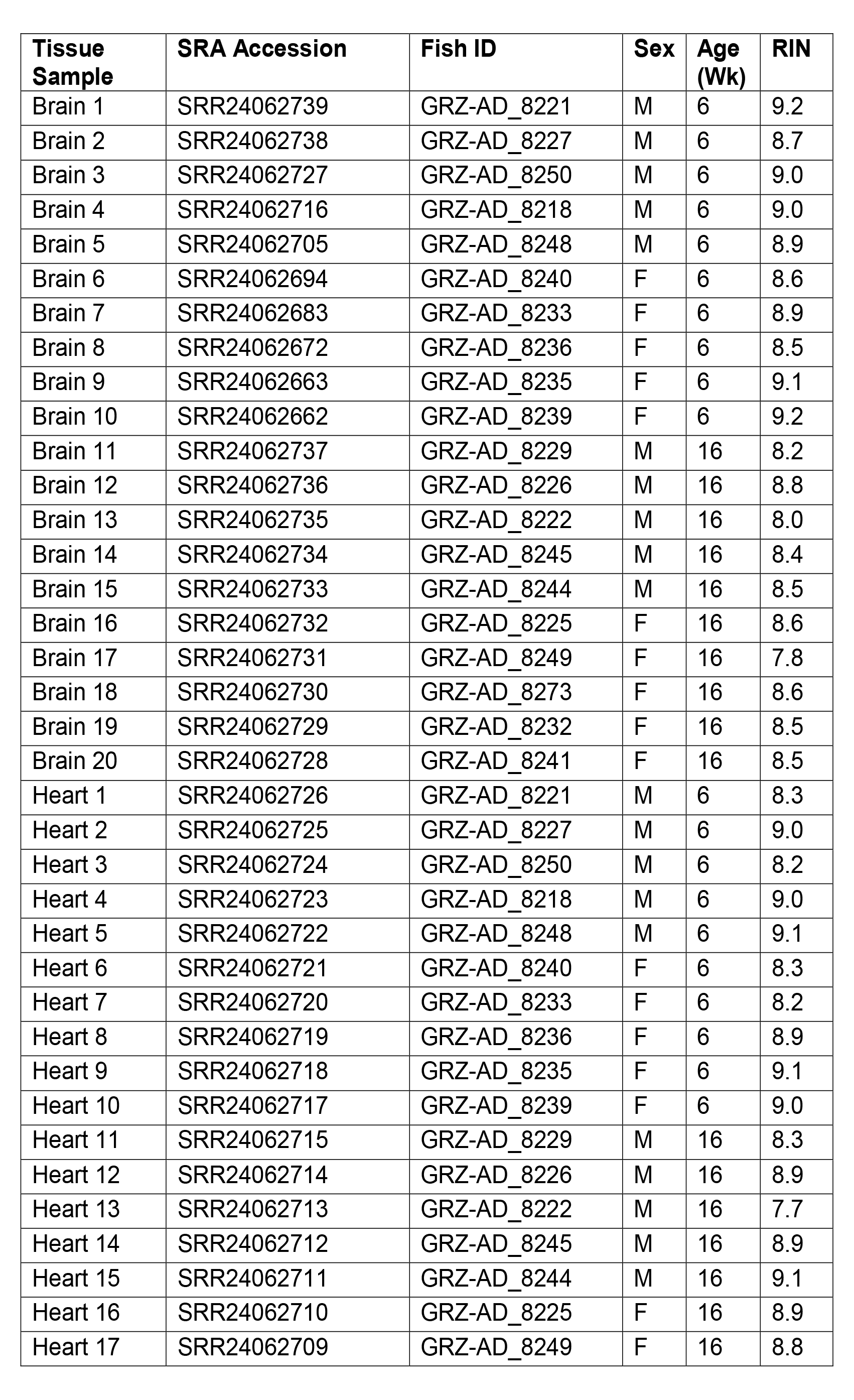
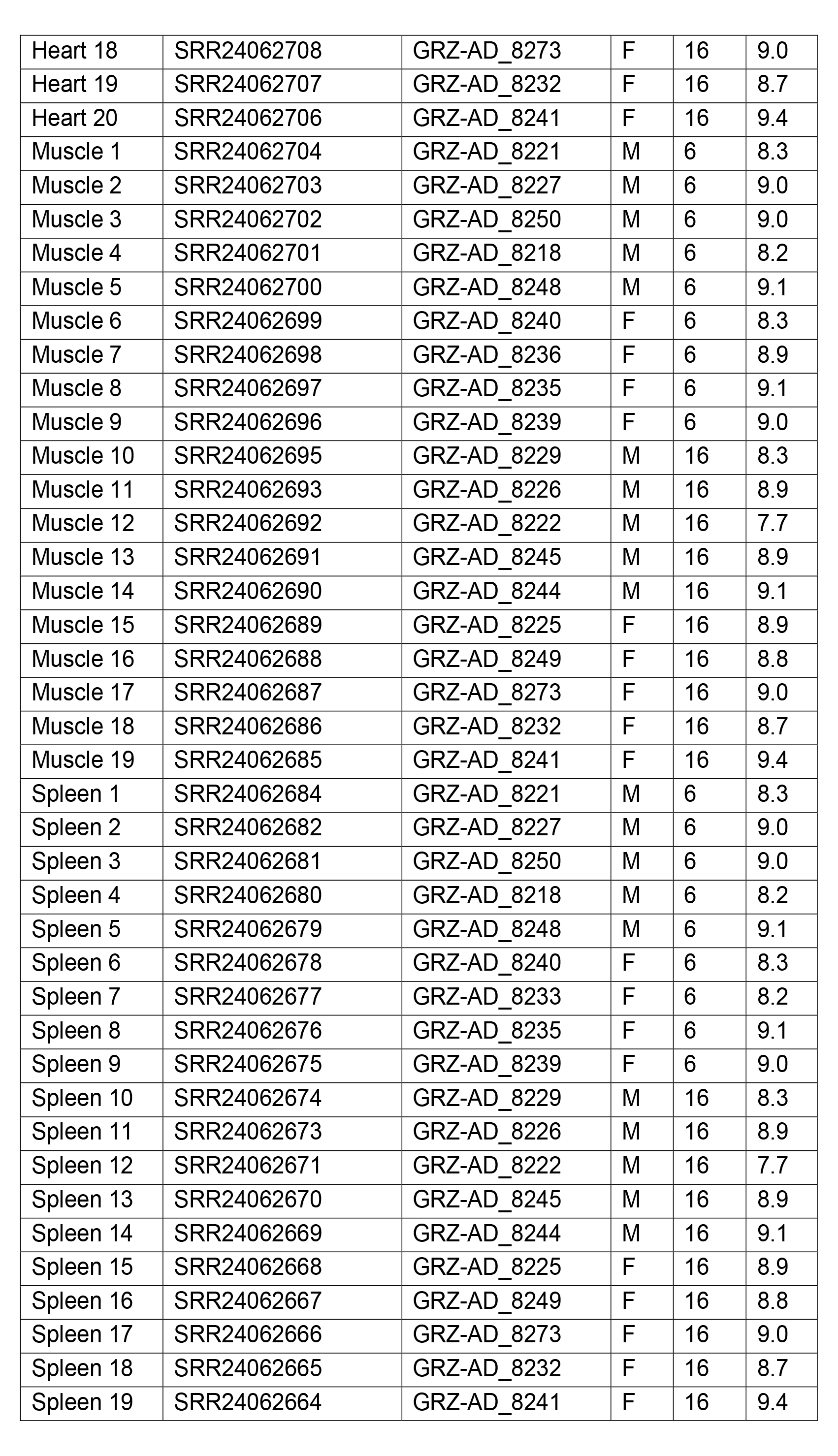
Sample information.

## Methods

### Fish Husbandry & Tissue Collection

Breeding, embryo collection, hatching, and fish husbandry followed standard protocols (Dodzian et al. 2018). Fish were reared in the fish facility at MPI age under §11TSchG animal housing license No. 576.1.36.6.G12/18 and euthanized (license MPIa_Anzeige_RB.16.005) by anesthetic overdose (600 mg/L MS222 in system water) administered by trained personnel. Fish were dissected to extract the brain, heart, liver, muscle and spleen which were immediately flash-frozen in liquid nitrogen and stored at −80˚C until use.

### RNA Isolation

For RNA isolation, frozen tissues (30-50mg) were placed in MP biomedicals lysis matrix D tubes (CAT#6913500) filled with 1mL of Trizol reagent (Thermo-Fisher), then homogenized using Benchmark BeadBug 6. Total RNA was purified using Direct-zol RNA Miniprep Plus Kit (Zymo cat# R2072) following the manufacturer’s instructions. RNA quality was assessed using high sensitivity RNA screen tapes (Agilent cat# 5067-5579, 5067-5580) on Agilent Tapestation 4200 to obtain the RNA Integrity Number (RIN). Samples with a RIN score of <4 were discarded, which excluded 8/20 liver samples including all old male samples. Due to the high number of samples that did not pass QC, which would compromise our ability to measure some of the biological groups, we chose not to proceed with liver RNA-seq library preparation.

### RNA-Seq library preparation and sequencing

We used 40ng of total RNA, which was subjected to ribosomal-RNA depletion using the RiboGone™ - Mammalian kit (Clontech cat# 634847) according to the manufacturer’s protocol. Strand specific RNA-seq libraries were then constructed using the SMARTer Stranded RNA-seq Kit (Clontech), according to the manufacturer’s protocol. Libraries were quality controlled using high sensitivity D1000 screen tapes (Agilent cat# 5067-5585, 5067-5603) on Agilent Tapestation 4200 before multiplexing the libraries for sequencing. Some samples were lost at this stage, as no library could be recovered, *i.e*. muscle sample 7 (young female) and spleen sample 8 (young female). Libraries that passed all QC steps were sequenced as paired-end 150-bp reads on the HiSeq X Ten platform at Novogene Corporation (USA).

### Bioinformatic Analysis

#### Adapter trimming and quality control

Raw reads were trimmed of adapters and low-quality reads were filtered using fastp version 0.23.2 with parameters “--failed_out fail_reads.out --detected_adapter_for_pe”. Raw reads and filtered reads were then quality-checked with Fastqc version 0.11.9 under default parameters. Multiqc version 1.15 was used to summarize the Fastqc reports.

#### Mapping and counting reads

Filtered reads were mapped to killifish reference genome (GCA_014300015.1) ^32^ that was softmasked with RepeatMasker version 4.1.2-p1^33^ with *Nothobranchius furzeri* TE sequences obtained from FishTEDB^34^ (as described in {Teefy, 2023 #3}), using STAR version 2.7.0e^35^ with parameters “--outFilterMultimapNmax 200 --outFilterIntronMotifs RemoveNoncanonicalUnannotated --alignEndsProtrude 10 ConcordantPair -- limitGenomeGenerateRAM 60000000000 --outSAMtype BAM SortedByCoordinate”. Multiqc version 1.15 was used to summarize the alignment reports generated by STAR. Gene and TE count matrices were generated against killifish reference gene annotation and the TE annotation using TEtranscripts version 2.2.1^36^ with parameter “--sortedByPos”.

To determine the ratio of reads mapped to introns and exons, we used featureCounts version 2.0.4^37^ to summarize the number of reads mapped to the exon level and gene level with the killifish reference gene annotation, respectively. The number of intronic reads was determined by subtracting the sum of exonic reads from the sum of reads mapped to gene features^38^.

#### Transposable element read ratio

To determine the ratio of reads contributed by TE regions, the TEtranscripts summarized count matrices were imported into R version 4.3.0^39^. The sum of reads mapped to TE features was divided by the total sum of reads in each tissue samples respectively. Non-parametric Mann-Whitney rank test was used to determine whether there was a statistically significant difference in TE ratio grouped by sex and age with ggpubr version 0.6.0^40^ and false discovery rate was reported to correct for multiple testing.

#### Differential gene expression analysis & transcriptional read correlation

The TEtranscripts summarized count matrices were imported into R version 4.3.0 and differential gene expression analysis was conducted using DESeq2 version 1.40.1^41^ with sex and age as modeling variables. Normalized count matrices, variance-stabilized count matrices and differential gene expression result matrices were generated. Transcriptome-wide correlation of reads mapped to gene and TE features was determined by assessing the pair-wise Spearman rank correlation between each sample pair. We also used principal component analysis on the variance-stabilized count matrices to determine the overall separation of samples across tissue types, as a function of age and sex.

#### Variance partition analysis

To determine the amount of variance that could be explained by sex and age, the variance-stabilized count matrices were first split into TE and canonical gene count matrix. R package variancePartition version 1.30.0^42^ was used to determine the amount of variance explained by sex and age in TE and canonical gene count matrices respectively.

#### Gene ontology analysis

To determine the biological pathways that were significantly altered in aging and pathways that were implicated in sex dimorphism, we performed GSEA (gene set enrichment analysis) GO analysis. As described in Teefy *et al*., turquoise killifish protein sequences from GCA_014300015.1 were aligned to the Ensembl release 104 human protein database using BLASTP (NCBI BLAST version 2.13.0). The top human protein sequence for each turquoise killifish hit was retained using a minimal E-value cutoff of 10^-3^ and used for GSEA. The results of the differential gene expression analysis with respect to sex and age were used as inputs for GSEA for each tissue. Killifish reference gene annotations were substituted with human homolog when possible and genes without human homologs that were able to pass the BLASTP filter were discarded. When multiple genes map to the same human homolog, the log-two-fold change were averaged. Genes were then sorted in a decreasing order with respect to the log-two-fold change. GSEA was performed using the R package clusterProfiler version 4.8.1^43^ and human gene ontology database org.Hs.eg.db version 3.17.0^44^. GSEA was run using a minimum gene set of 25 terms and a maximum gene set of 5,000 terms with a significance cutoff of FDR <5%.

### Data Records

Sequencing data was submitted to the Sequence Read Archive and is accessible through BioProject PRJNA952180: Transcriptional profiling of aging tissues from African turquoise killifish.

## Technical Validation

### Experimental design and quality control

We generated ribosomal RNA-depleted RNA-seq libraries from the brain, heart, muscle, and spleen from young (6-weeks-old) and old (16-weeks-old) male and female GRZ strain turquoise killifish starting from 5 fish per biological group (Figure 1A, Table 1). Importantly, each euthanized fish contributed all profiled tissues to minimize the number of subjects, as well as ultimately to potentially identify transcriptional signatures common to particular subjects across multiple tissues (*i.e*. brain, heart, muscle, and spleen samples from individual GRZ-AD_8240; Table 1). Due to library construction failure, one young female muscle sample and one young female spleen sample were not sequenced (Table 1).

We began to assess library quality by analyzing the number of reads in each library (Figure 2A, Table 2). Each library had roughly the same number of counts with no systemic bias towards any groups. Of note, the brain libraries consistently had the fewest total reads, although read counts were comparable across brain libraries. Next, we performed FastQC analysis using the MultiQC tool on each RNA-seq library to determine the mean quality scores for each sample (Figure 2B). Quality scores for each library consistently had a Phred score >= 33 for the length of the read thereby, showing we generated high-quality RNA-seq libraries.

**Figure 2.**
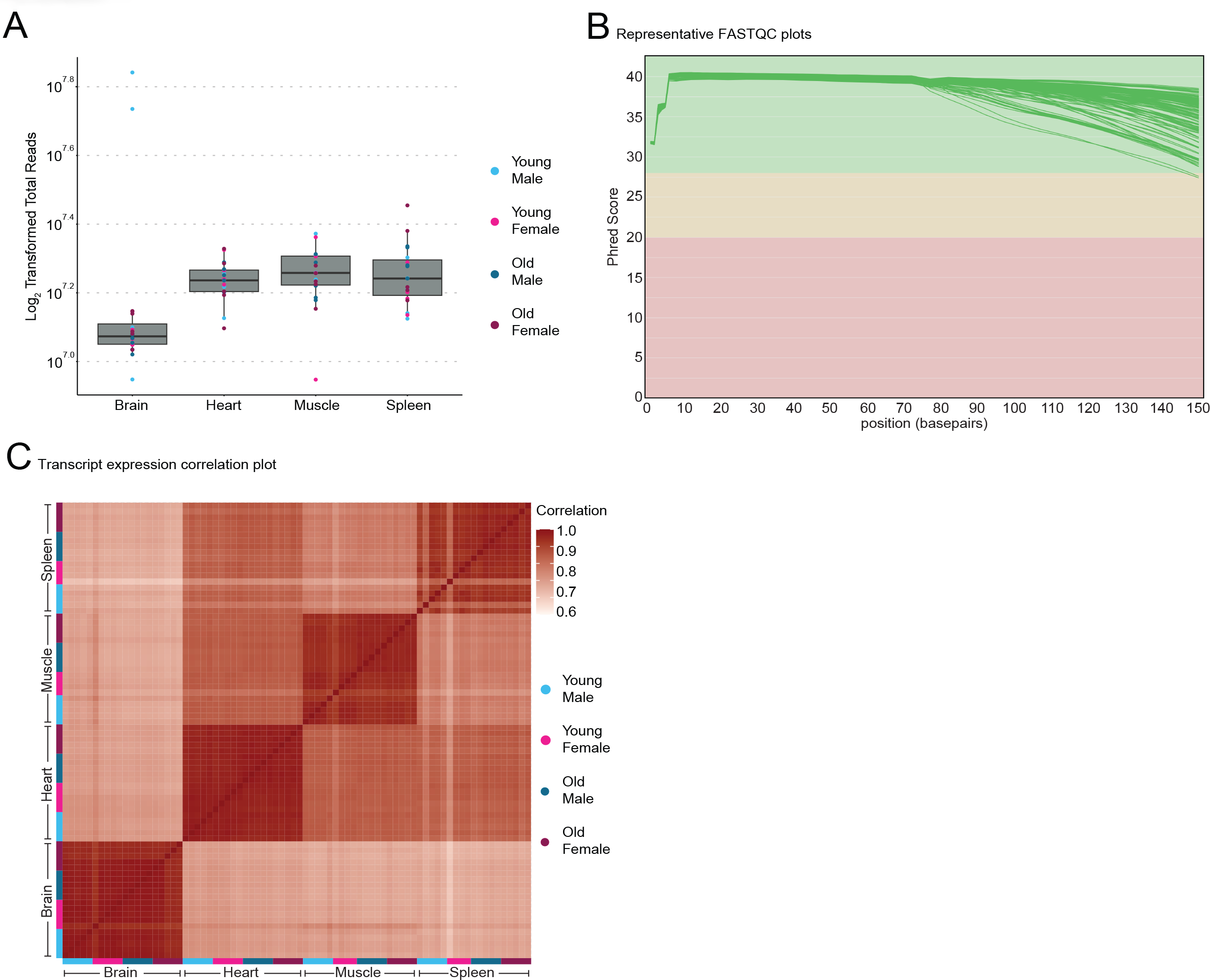
Quality control metrics for RNA-seq libraries of African turquoise killifish tissues. (A) Boxplot of raw log_2_ counts from each library. Count distributions are similar between all replicates indicating unbiased sample preparation. (B) Representative output from MulitQC showing the Phred score for each library across the length of the reads. Each sample shows high read quality with a Phred score that is >33 for the majority of the read length. (C) Correlation plot of all RNA-seq libraries. Libraries common to each tissue tend to correlate extremely tightly. The brain appears have the most distinct transcriptional profile of all tissues in our dataset.

**Table 2.**
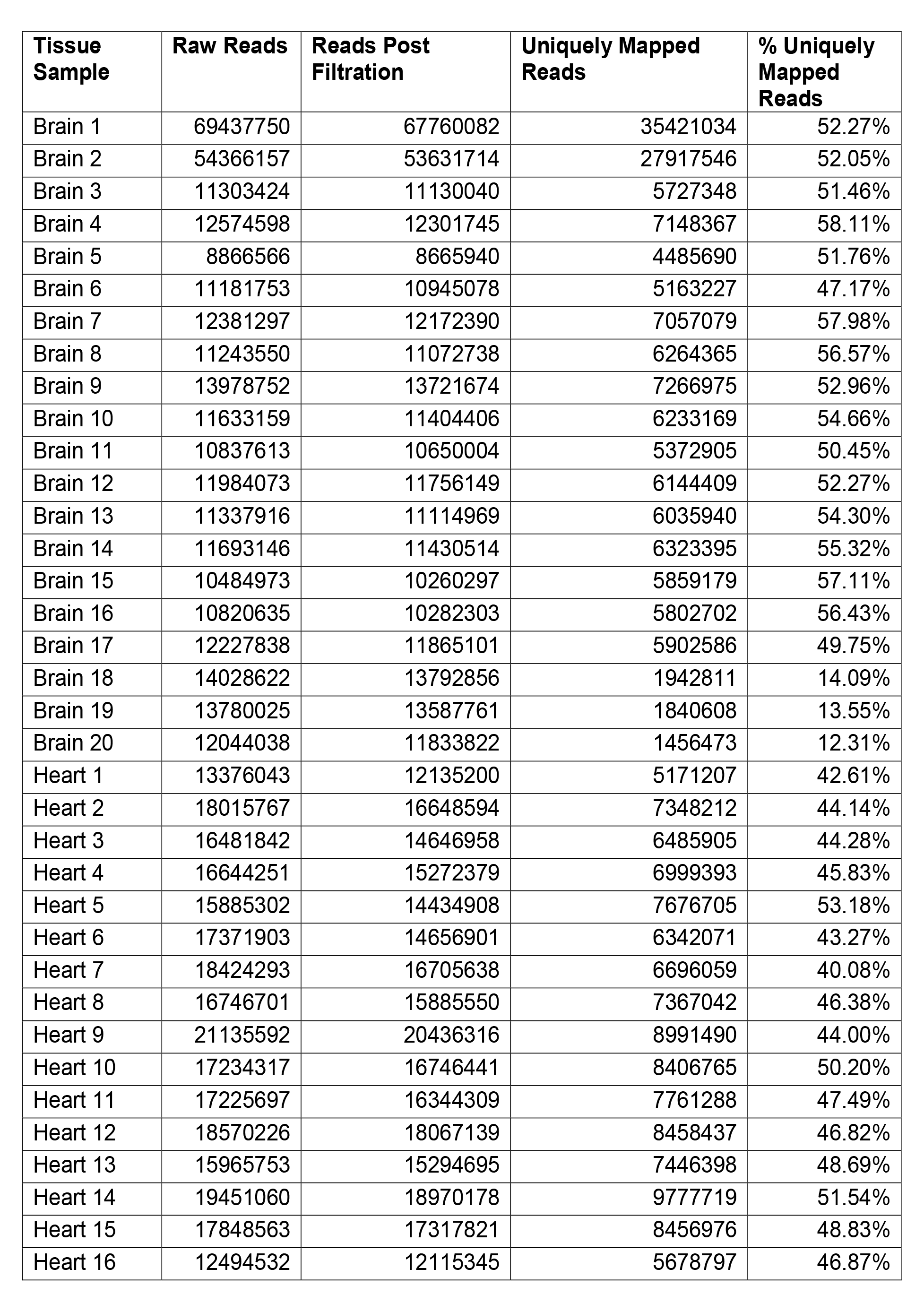
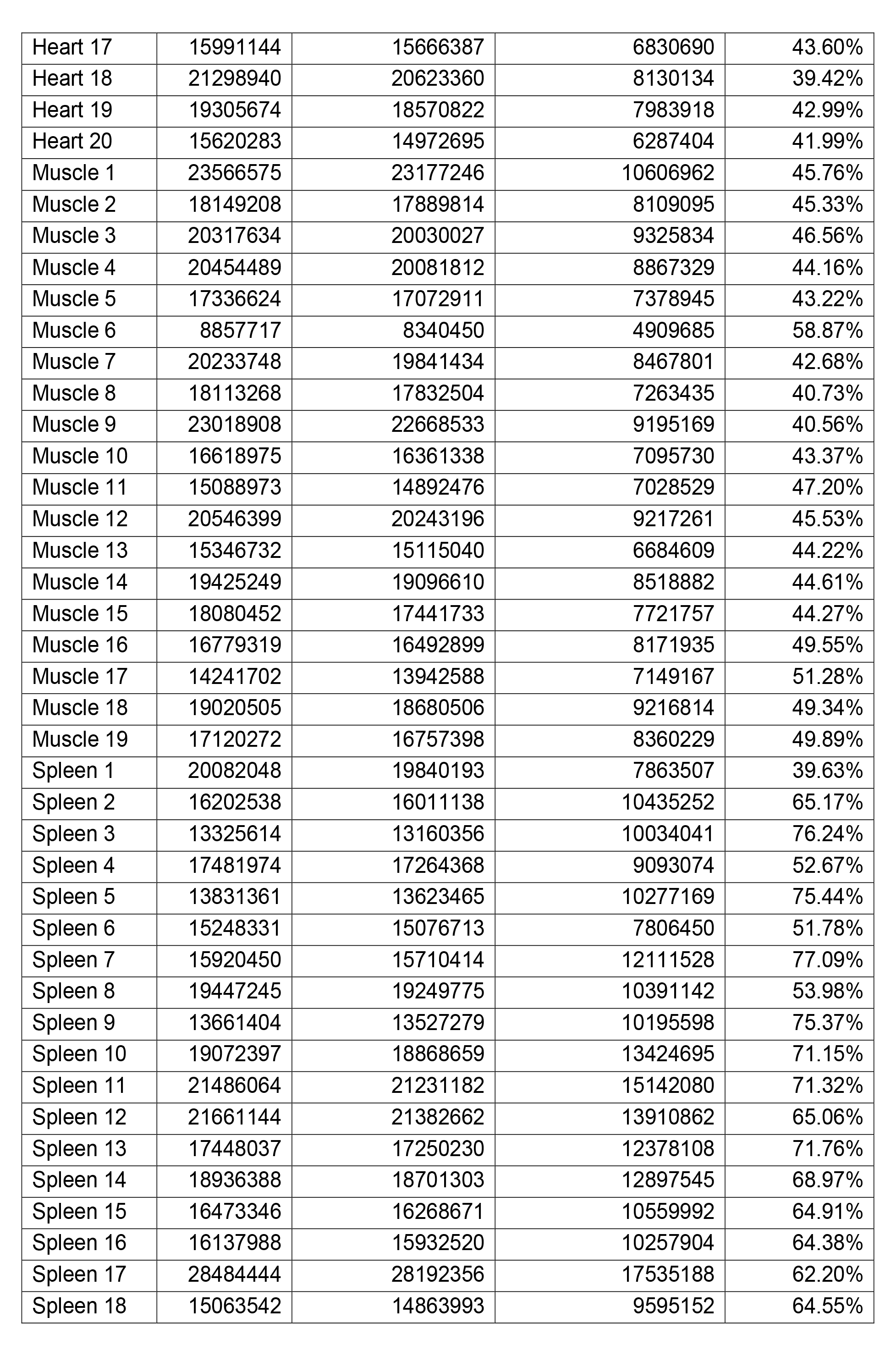
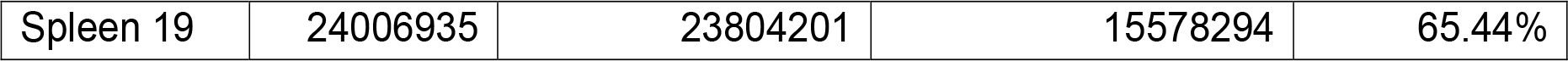
Read information.

| **Tissue Sample** | **Raw Reads** | **Reads Post Filtration** | **Uniquely Mapped Reads** | **% Uniquely Mapped Reads** |
| --- | --- | --- | --- | --- |
| Brain 1 | 69437750 | 67760082 | 35421034 | 52.27% |
| Brain 2 | 54366157 | 53631714 | 27917546 | 52.05% |
| Brain 3 | 11303424 | 11130040 | 5727348 | 51.46% |
| Brain 4 | 12574598 | 12301745 | 7148367 | 58.11% |
| Brain 5 | 8866566 | 8665940 | 4485690 | 51.76% |
| Brain 6 | 11181753 | 10945078 | 5163227 | 47.17% |
| Brain 7 | 12381297 | 12172390 | 7057079 | 57.98% |
| Brain 8 | 11243550 | 11072738 | 6264365 | 56.57% |
| Brain 9 | 13978752 | 13721674 | 7266975 | 52.96% |
| Brain 10 | 11633159 | 11404406 | 6233169 | 54.66% |
| Brain 11 | 10837613 | 10650004 | 5372905 | 50.45% |
| Brain 12 | 11984073 | 11756149 | 6144409 | 52.27% |
| Brain 13 | 11337916 | 11114969 | 6035940 | 54.30% |
| Brain 14 | 11693146 | 11430514 | 6323395 | 55.32% |
| Brain 15 | 10484973 | 10260297 | 5859179 | 57.11% |
| Brain 16 | 10820635 | 10282303 | 5802702 | 56.43% |
| Brain 17 | 12227838 | 11865101 | 5902586 | 49.75% |
| Brain 18 | 14028622 | 13792856 | 1942811 | 14.09% |
| Brain 19 | 13780025 | 13587761 | 1840608 | 13.55% |
| Brain 20 | 12044038 | 11833822 | 1456473 | 12.31% |
| Heart 1 | 13376043 | 12135200 | 5171207 | 42.61% |
| Heart 2 | 18015767 | 16648594 | 7348212 | 44.14% |
| Heart 3 | 16481842 | 14646958 | 6485905 | 44.28% |
| Heart 4 | 16644251 | 15272379 | 6999393 | 45.83% |
| Heart 5 | 15885302 | 14434908 | 7676705 | 53.18% |
| Heart 6 | 17371903 | 14656901 | 6342071 | 43.27% |
| Heart 7 | 18424293 | 16705638 | 6696059 | 40.08% |
| Heart 8 | 16746701 | 15885550 | 7367042 | 46.38% |
| Heart 9 | 21135592 | 20436316 | 8991490 | 44.00% |
| Heart 10 | 17234317 | 16746441 | 8406765 | 50.20% |
| Heart 11 | 17225697 | 16344309 | 7761288 | 47.49% |
| Heart 12 | 18570226 | 18067139 | 8458437 | 46.82% |
| Heart 13 | 15965753 | 15294695 | 7446398 | 48.69% |
| Heart 14 | 19451060 | 18970178 | 9777719 | 51.54% |
| Heart 15 | 17848563 | 17317821 | 8456976 | 48.83% |
| Heart 16 | 12494532 | 12115345 | 5678797 | 46.87% |
| Heart 17 | 15991144 | 15666387 | 6830690 | 43.60% |
| Heart 18 | 21298940 | 20623360 | 8130134 | 39.42% |
| Heart 19 | 19305674 | 18570822 | 7983918 | 42.99% |
| Heart 20 | 15620283 | 14972695 | 6287404 | 41.99% |
| Muscle 1 | 23566575 | 23177246 | 10606962 | 45.76% |
| Muscle 2 | 18149208 | 17889814 | 8109095 | 45.33% |
| Muscle 3 | 20317634 | 20030027 | 9325834 | 46.56% |
| Muscle 4 | 20454489 | 20081812 | 8867329 | 44.16% |
| Muscle 5 | 17336624 | 17072911 | 7378945 | 43.22% |
| Muscle 6 | 8857717 | 8340450 | 4909685 | 58.87% |
| Muscle 7 | 20233748 | 19841434 | 8467801 | 42.68% |
| Muscle 8 | 18113268 | 17832504 | 7263435 | 40.73% |
| Muscle 9 | 23018908 | 22668533 | 9195169 | 40.56% |
| Muscle 10 | 16618975 | 16361338 | 7095730 | 43.37% |
| Muscle 11 | 15088973 | 14892476 | 7028529 | 47.20% |
| Muscle 12 | 20546399 | 20243196 | 9217261 | 45.53% |
| Muscle 13 | 15346732 | 15115040 | 6684609 | 44.22% |
| Muscle 14 | 19425249 | 19096610 | 8518882 | 44.61% |
| Muscle 15 | 18080452 | 17441733 | 7721757 | 44.27% |
| Muscle 16 | 16779319 | 16492899 | 8171935 | 49.55% |
| Muscle 17 | 14241702 | 13942588 | 7149167 | 51.28% |
| Muscle 18 | 19020505 | 18680506 | 9216814 | 49.34% |
| Muscle 19 | 17120272 | 16757398 | 8360229 | 49.89% |
| Spleen 1 | 20082048 | 19840193 | 7863507 | 39.63% |
| Spleen 2 | 16202538 | 16011138 | 10435252 | 65.17% |
| Spleen 3 | 13325614 | 13160356 | 10034041 | 76.24% |
| Spleen 4 | 17481974 | 17264368 | 9093074 | 52.67% |
| Spleen 5 | 13831361 | 13623465 | 10277169 | 75.44% |
| Spleen 6 | 15248331 | 15076713 | 7806450 | 51.78% |
| Spleen 7 | 15920450 | 15710414 | 12111528 | 77.09% |
| Spleen 8 | 19447245 | 19249775 | 10391142 | 53.98% |
| Spleen 9 | 13661404 | 13527279 | 10195598 | 75.37% |
| Spleen 10 | 19072397 | 18868659 | 13424695 | 71.15% |
| Spleen 11 | 21486064 | 21231182 | 15142080 | 71.32% |
| Spleen 12 | 21661144 | 21382662 | 13910862 | 65.06% |
| Spleen 13 | 17448037 | 17250230 | 12378108 | 71.76% |
| Spleen 14 | 18936388 | 18701303 | 12897545 | 68.97% |
| Spleen 15 | 16473346 | 16268671 | 10559992 | 64.91% |
| Spleen 16 | 16137988 | 15932520 | 10257904 | 64.38% |
| Spleen 17 | 28484444 | 28192356 | 17535188 | 62.20% |
| Spleen 18 | 15063542 | 14863993 | 9595152 | 64.55% |
| Spleen 19 | 24006935 | 23804201 | 15578294 | 65.44% |

After confirming high-quality libraries, we mapped each RNA-seq library to a recently published killifish genome version^32^ that was softmasked with turquoise killifish TE sequences from FishTEDB^34^. First, we measured the intronic/exonic read ratio using featureCounts and observed that roughly half of all reads were intronic and half were exonic. The brain and spleen had the highest ratio of intronic reads while the heart and muscle had the lowest (Table 3).

**Table 3.** Read origin.

| Sample | Exonic Reads | Intronic Reads | Total Reads | % Exonic Reads | % Intronic Reads |
| --- | --- | --- | --- | --- | --- |
| Brain 1 | 14908766 | 14524642 | 29433408 | 50.65% | 49.35% |
| Brain 2 | 11605211 | 11231021 | 22836232 | 50.82% | 49.18% |
| Brain 3 | 2459842 | 2212442 | 4672284 | 52.65% | 47.35% |
| Brain 4 | 3025051 | 3046561 | 6071612 | 49.82% | 50.18% |
| Brain 5 | 1859843 | 1882245 | 3742088 | 49.70% | 50.30% |
| Brain 6 | 2250150 | 2060813 | 4310963 | 52.20% | 47.80% |
| Brain 7 | 2860757 | 2990514 | 5851271 | 48.89% | 51.11% |
| Brain 8 | 2552918 | 2614628 | 5167546 | 49.40% | 50.60% |
| Brain 9 | 3019711 | 3044151 | 6063862 | 49.80% | 50.20% |
| Brain 10 | 2533074 | 2653154 | 5186228 | 48.84% | 51.16% |
| Brain 11 | 2084136 | 2265343 | 4349479 | 47.92% | 52.08% |
| Brain 12 | 2371209 | 2615001 | 4986210 | 47.56% | 52.44% |
| Brain 13 | 2295764 | 2592469 | 4888233 | 46.97% | 53.03% |
| Brain 14 | 2527948 | 2703993 | 5231941 | 48.32% | 51.68% |
| Brain 15 | 2221703 | 2595674 | 4817377 | 46.12% | 53.88% |
| Brain 16 | 2144889 | 2668715 | 4813604 | 44.56% | 55.44% |
| Brain 17 | 2159375 | 2668819 | 4828194 | 44.72% | 55.28% |
| Brain 18 | 642867 | 725168 | 1368035 | 46.99% | 53.01% |
| Brain 19 | 638294 | 684914 | 1323208 | 48.24% | 51.76% |
| Brain 20 | 511203 | 512595 | 1023798 | 49.93% | 50.07% |
| Heart 1 | 2504484 | 1653732 | 4158216 | 60.23% | 39.77% |
| Heart 2 | 3618755 | 2423548 | 6042303 | 59.89% | 40.11% |
| Heart 3 | 3198633 | 2048401 | 5247034 | 60.96% | 39.04% |
| Heart 4 | 3402551 | 2280051 | 5682602 | 59.88% | 40.12% |
| Heart 5 | 3624183 | 2787411 | 6411594 | 56.53% | 43.47% |
| Heart 6 | 2950664 | 2255367 | 5206031 | 56.68% | 43.32% |
| Heart 7 | 3176159 | 2246760 | 5422919 | 58.57% | 41.43% |
| Heart 8 | 3611783 | 2516323 | 6128106 | 58.94% | 41.06% |
| Heart 9 | 4362438 | 2999203 | 7361641 | 59.26% | 40.74% |
| Heart 10 | 3906128 | 2803345 | 6709473 | 58.22% | 41.78% |
| Heart 11 | 3589614 | 2361223 | 5950837 | 60.32% | 39.68% |
| Heart 12 | 3909413 | 2658456 | 6567869 | 59.52% | 40.48% |
| Heart 13 | 3364117 | 2435300 | 5799417 | 58.01% | 41.99% |
| Heart 14 | 4541783 | 3130903 | 7672686 | 59.19% | 40.81% |
| Heart 15 | 3960859 | 2637727 | 6598586 | 60.03% | 39.97% |
| Heart 16 | 2754188 | 1787142 | 4541330 | 60.65% | 39.35% |
| Heart 17 | 3220970 | 2084119 | 5305089 | 60.71% | 39.29% |
| Heart 18 | 4034668 | 2609811 | 6644479 | 60.72% | 39.28% |
| Heart 19 | 3669026 | 2713662 | 6382688 | 57.48% | 42.52% |
| Heart 20 | 3029837 | 2160539 | 5190376 | 58.37% | 41.63% |
| Muscle 1 | 7807559 | 2564152 | 10371711 | 75.28% | 24.72% |
| Muscle 2 | 5965916 | 1893177 | 7859093 | 75.91% | 24.09% |
| Muscle 3 | 6701665 | 2036250 | 8737915 | 76.70% | 23.30% |
| Muscle 4 | 6669478 | 2069602 | 8739080 | 76.32% | 23.68% |
| Muscle 5 | 4912246 | 1903882 | 6816128 | 72.07% | 27.93% |
| Muscle 6 | 3541551 | 1276645 | 4818196 | 73.50% | 26.50% |
| Muscle 7 | 6062051 | 1852763 | 7914814 | 76.59% | 23.41% |
| Muscle 8 | 5184648 | 1718353 | 6903001 | 75.11% | 24.89% |
| Muscle 9 | 6506879 | 2179090 | 8685969 | 74.91% | 25.09% |
| Muscle 10 | 4370183 | 1704847 | 6075030 | 71.94% | 28.06% |
| Muscle 11 | 4583912 | 1722128 | 6306040 | 72.69% | 27.31% |
| Muscle 12 | 5955858 | 2129553 | 8085411 | 73.66% | 26.34% |
| Muscle 13 | 4620295 | 1587798 | 6208093 | 74.42% | 25.58% |
| Muscle 14 | 5555667 | 1867962 | 7423629 | 74.84% | 25.16% |
| Muscle 15 | 5363929 | 1871312 | 7235241 | 74.14% | 25.86% |
| Muscle 16 | 5679905 | 2048191 | 7728096 | 73.50% | 26.50% |
| Muscle 17 | 5281032 | 1675725 | 6956757 | 75.91% | 24.09% |
| Muscle 18 | 6467530 | 2129778 | 8597308 | 75.23% | 24.77% |
| Muscle 19 | 5815008 | 1991870 | 7806878 | 74.49% | 25.51% |
| Spleen 1 | 4334557 | 2639241 | 6973798 | 62.15% | 37.85% |
| Spleen 2 | 5843374 | 3410146 | 9253520 | 63.15% | 36.85% |
| Spleen 3 | 4753534 | 4274154 | 9027688 | 52.66% | 47.34% |
| Spleen 4 | 4907508 | 3432296 | 8339804 | 58.84% | 41.16% |
| Spleen 5 | 4821133 | 4489991 | 9311124 | 51.78% | 48.22% |
| Spleen 6 | 3860571 | 3240579 | 7101150 | 54.37% | 45.63% |
| Spleen 7 | 5613011 | 5396055 | 11009066 | 50.99% | 49.01% |
| Spleen 8 | 5699582 | 3757365 | 9456947 | 60.27% | 39.73% |
| Spleen 9 | 4786156 | 4463578 | 9249734 | 51.74% | 48.26% |
| Spleen 10 | 6453914 | 5638011 | 12091925 | 53.37% | 46.63% |
| Spleen 11 | 7558220 | 6235012 | 13793232 | 54.80% | 45.20% |
| Spleen 12 | 6993332 | 5508069 | 12501401 | 55.94% | 44.06% |
| Spleen 13 | 5759573 | 5198553 | 10958126 | 52.56% | 47.44% |
| Spleen 14 | 6332649 | 5297385 | 11630034 | 54.45% | 45.55% |
| Spleen 15 | 5097488 | 4403099 | 9500587 | 53.65% | 46.35% |
| Spleen 16 | 5105004 | 4209854 | 9314858 | 54.80% | 45.20% |
| Spleen 17 | 8749232 | 7240568 | 15989800 | 54.72% | 45.28% |
| Spleen 18 | 4584090 | 4046504 | 8630594 | 53.11% | 46.89% |
| Spleen 19 | 7752928 | 6353132 | 14106060 | 54.96% | 45.04% |

Next, to capture both gene and TE counts, we generated read counts for genes and TEs using TETranscripts, as in Teefy *et al*.^17^. After generating count matrices consisting of gene and TE counts, we normalized reads in DESeq2 and created transcript expression correlation maps between libraries (Figure 2C). We found that samples clustered tightly by tissue, consistent with strong expected tissue-specific transcript expression. To assess transcriptional similarity of various samples in each tissue, we performed principal component analysis (PCA) on each count matrices normalized with the Variance Stabilizing Transformation in DESeq2 (Figure 3A- D). In all tissues, transcript expression tended to segregate mostly by age, with a lesser secondary separation by sex. To quantify how much transcript expression variation in each tissue could be explained by age and sex, we used “variancePartition” (Figure 3E-F). Interestingly, in each tissue, age accounted for more variance in gene expression than sex for both genes (Figure 3E) and TEs (Figure 3F).

**Figure 3.**
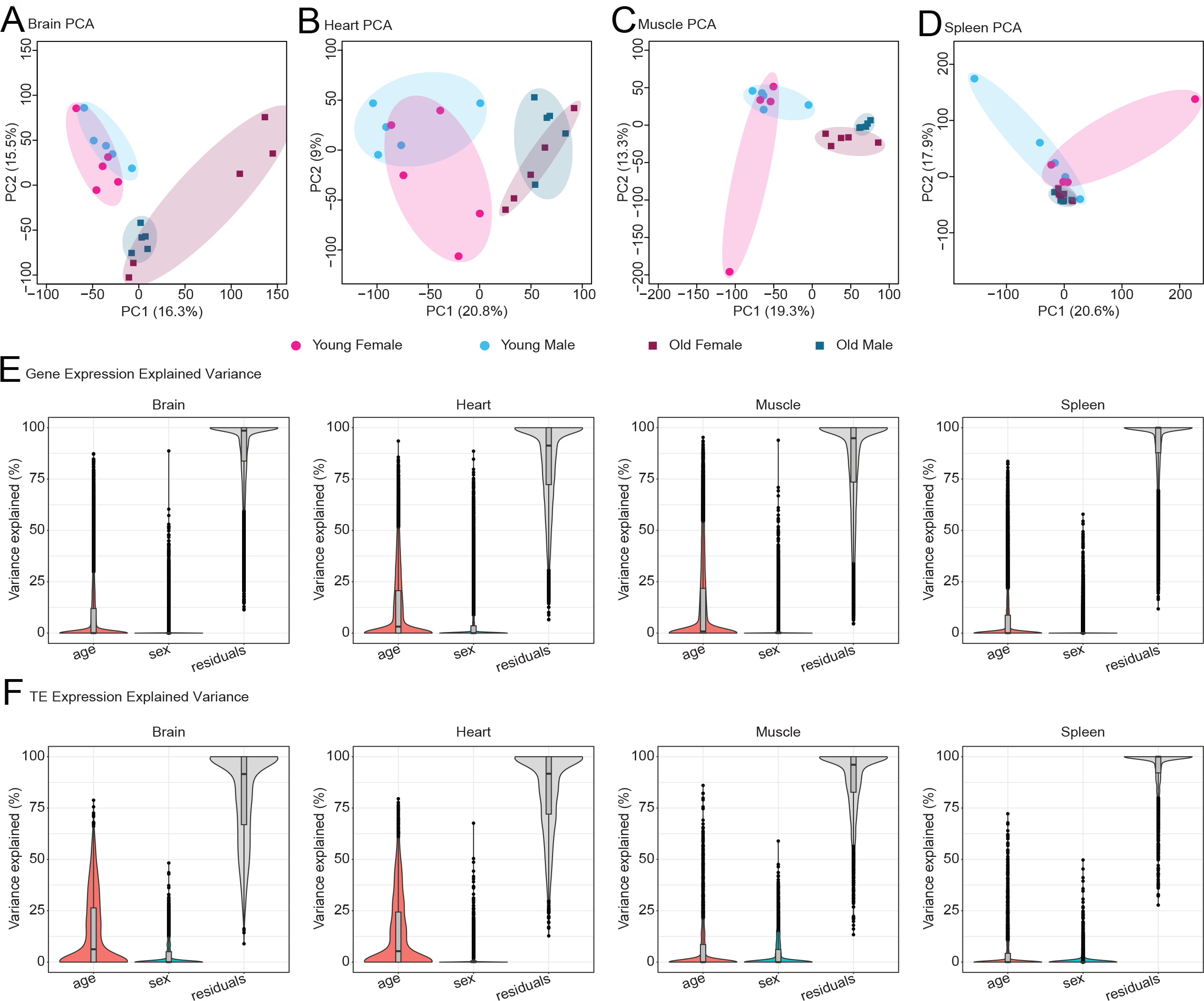
PCA plots of African turquoise killifish tissue transcriptomes as a function of aging and sex. (A-D) PCA plots of transcript expression highlighted by group (young female, young male, old female, old male) for (A) brain (B) heart (C) muscle (D) spleen. In each tissue besides the spleen, transcript expression segregates by age along PC1. In the spleen, samples still separate by age but primarily along PC2. (E) Variance in gene expression explained by age and sex in each tissue. In each tissue, age explains more of the variance in gene expression relative to sex. (F) Variance in TE expression explained by age and sex. In each tissue, age explains more of the variance in TE expression relative to sex.

### Differential transcription by age and sex

To assess the quality and useability of the dataset, we next performed differential gene expression analysis using DESeq2, starting by using only genes, and then with TEs (see below). We used a combined differential expression model with animal age and sex as modeling covariates. Using a significance threshold of FDR < 5%, we identified substantial age-related gene expression changes with 3611, 4910, 5077, and 2195 differentially expressed genes in brain, heart, muscle, and spleen, respectively (Figure 4A). In agreement with our PCA analysis, we find fewer genes with sex-dimorphic expression in each tissue with 0, 429, 30, and 13 differentially expressed genes between females and males in the brain, heart, muscle, and spleen, respectively (Figure 4B).

**Figure 4.**
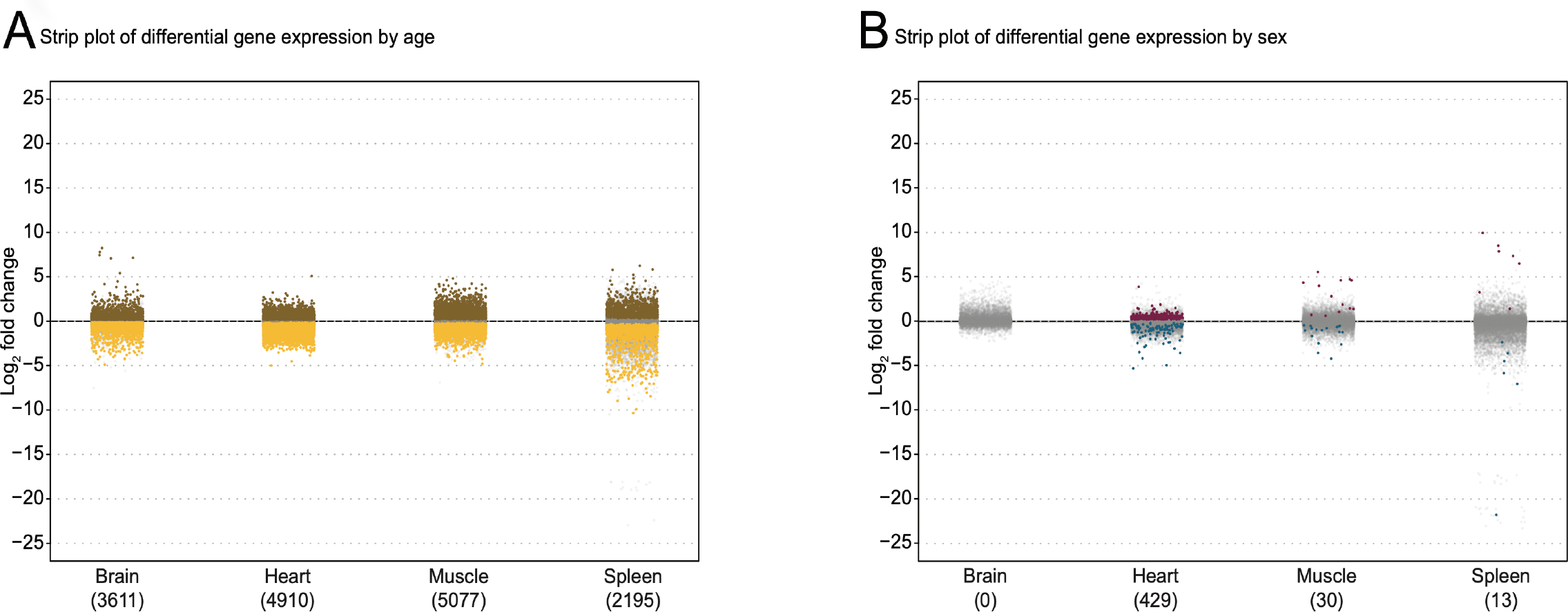
Differential gene expression analysis of African turquoise killifish tissue transcriptomes as a function of aging and sex. (A) Strip plot showing the number of differentially expressed genes by age (FDR < 5%) with the number of significantly differential genes in parentheses. In each tissue, thousands of genes are differentially expressed by age. The muscle was the tissue most affected by age at the transcriptional level with 5,077 differentially expressed genes. Brown denotes genes upregulated in old tissues, yellow denotes genes upregulated in young tissues, gray denotes non-significant differences in gene expression between ages. (B) Strip plot showing the number of differentially expressed genes by sex (FDR < 5%) with the number of significantly differential genes in parentheses. Fewer genes are differentially expressed by sex than age in all tissues assayed. The heart had the most sex-dimorphic gene expression with 429 genes differentially expressed by sex. Pink denotes genes upregulated in female tissues, blue denotes genes upregulated in male tissues, gray signifies genes with no significant expression differences between sexes.

Next, we analyzed the differential expression of TEs in these tissues. We found that as a percentage of mapped reads in each library, reads mapping to TEs ranges varied strongly by tissue, with ∼50% of all reads in brain libraries mapping to TEs and only < 20% of reads mapping to TEs in muscle libraries (Figure 5A). TEs were more differentially expressed by age rather than sex with 897, 706, 291, and 114 differentially expressed TEs in brain, heart, muscle, and spleen, respectively. Most tissues had an approximately equal proportion of up-and down-regulated TEs except the brain, which showed a strong bias for TE upregulation with age (Figure 5B). Like genes, TEs had more limited sex-dimorphic expression compared to age-related expression with only 15, 14, 0, and 1 differentially expressed TEs between sexes in brain, heart, muscle, and spleen, respectively (Figure 5C).

**Figure 5.**
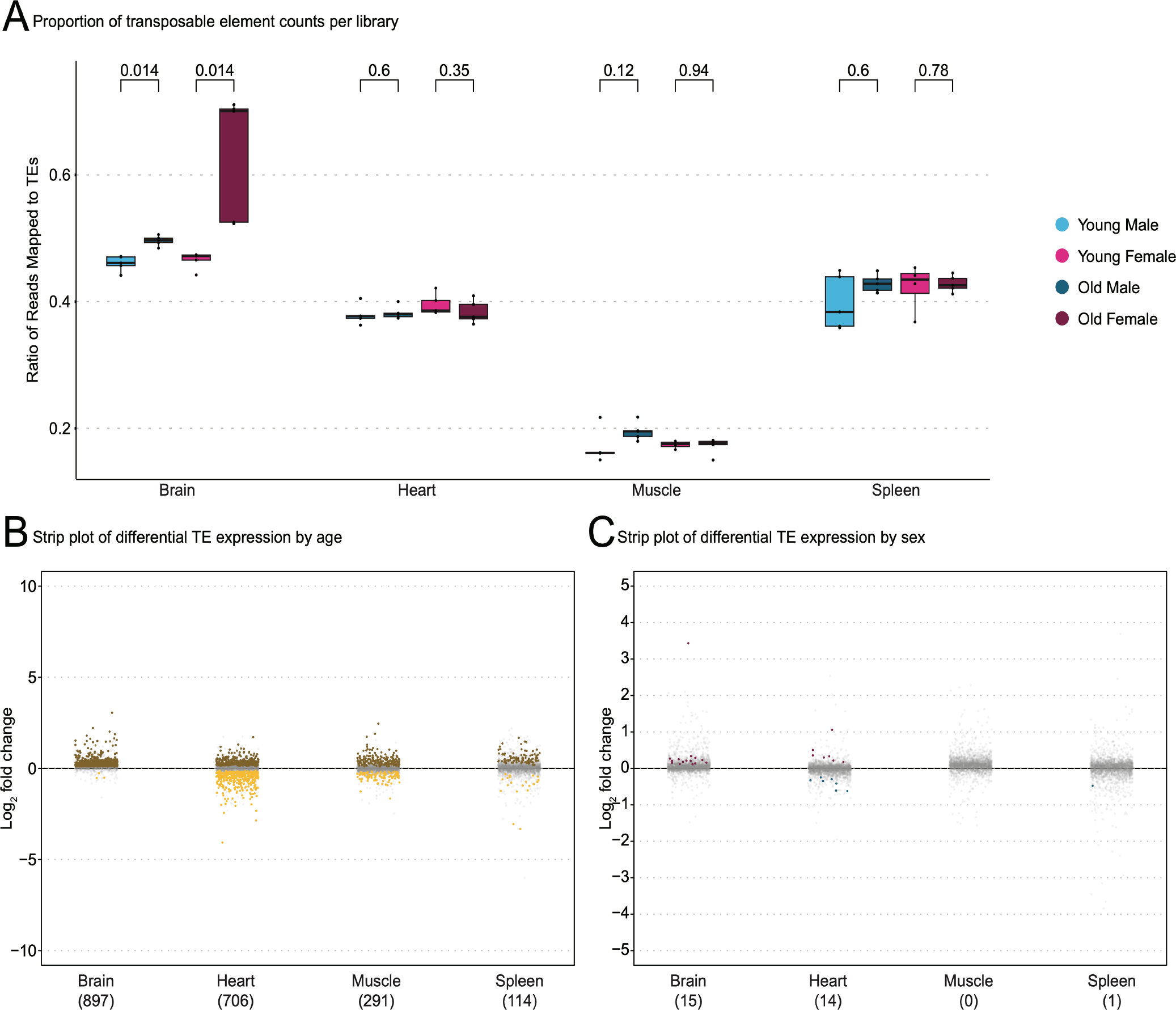
Transposable element quality control and differential expression analysis. (A) Boxplot depicting the relative proportions of counts attributed to genes and TEs in each library. The brain contains the most reads attributed to TEs by proportion with approximately half of all reads mapping to TEs while muscle has very few reads mapping to TEs. Significantly more reads map to TEs in old brains relative to young brains as measured by Wilcoxon test. Brown denotes TEs upregulated in old tissues, yellow denotes TEs upregulated in young tissues, gray denotes non-significant differences in gene expression between ages. (B) Strip plot of differentially expressed TEs by age (FDR < 5%) with the number of significantly differential TEs in parentheses. There is a substantial bias for TE upregulation in old brains. (C) Strip plot of differentially expressed TEs by sex (FDR < 5%) with the number of significantly differential TEs in parentheses. There are far fewer TEs differentially expressed by sex compared to age without an obvious bias towards any sex. Pink denotes TEs upregulated in female tissues, blue denotes TEs upregulated in male tissues, gray denotes non-significant differences in gene expression between sexes.

Lastly, we performed gene set enrichment analysis (GSEA) using gene ontology (GO) functional categories (using homology mapping from human annotations), to determine whether our dataset was amenable to this type of analysis. GO enrichment analysis was performed in each tissue, to determine enrichment as a function of age (Figure 6A) and as a function of sex (Figure 6B). As reported in previous aging ‘omic’ studies across animal taxa^2, 45^, at least one immune-related term was enriched in aged tissues compared to young tissues (Figure 6A), consistent with the notion of “inflamm-aging”. Importantly, young muscle also showed an enrichment of cell-cycle gene transcription, which may reflect more active or abundant muscle stem cells. All tissues displayed enough transcriptional sex-dimorphism to have at least 5 significantly enriched GO terms per sex, except for the female spleen, which only showed increased interferon production relative to the male spleen (Figure 6B).

**Figure 6.**
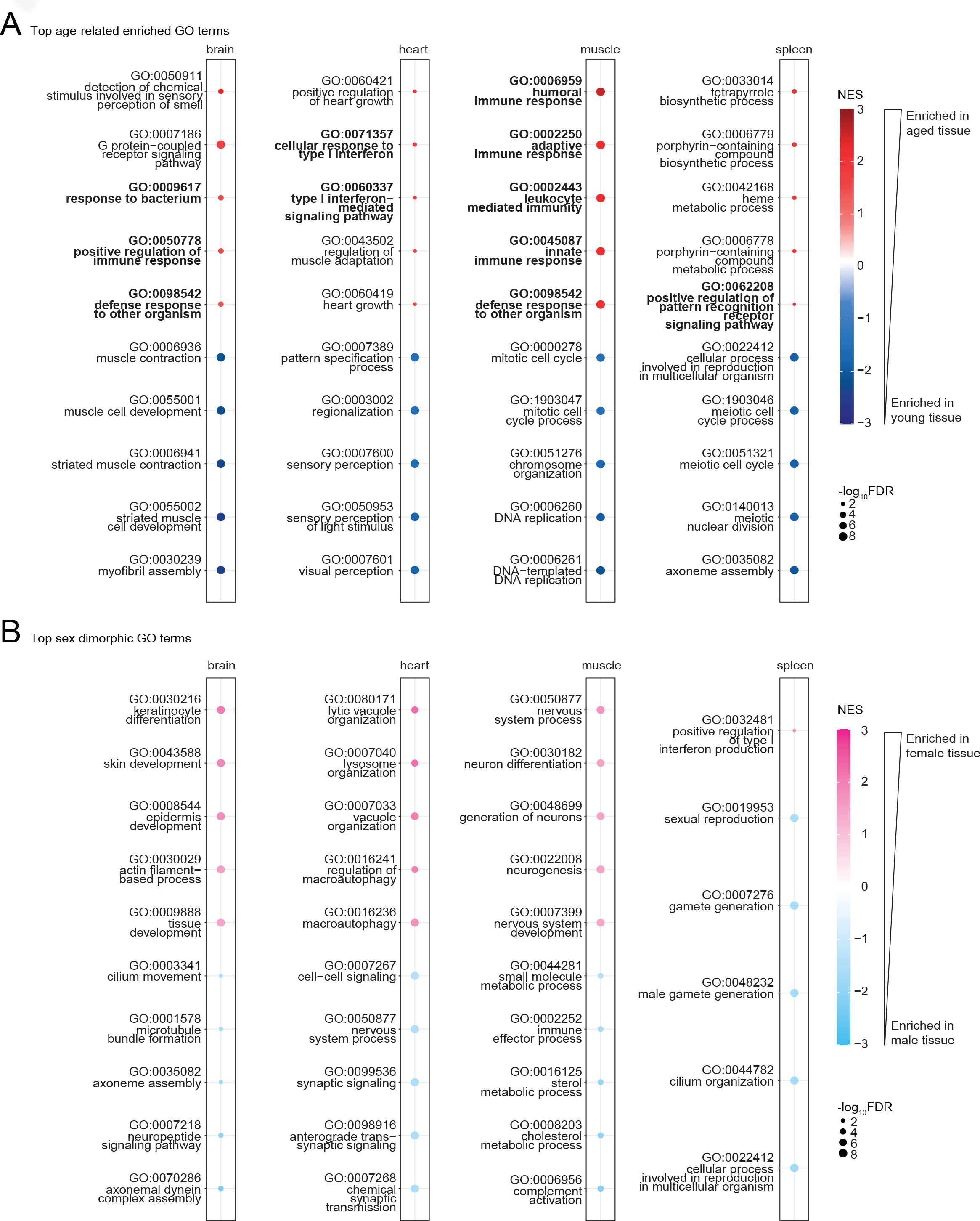
Gene Ontology enrichment analysis across African turquoise killifish tissues as a function of sex and age. (A) Top GO terms enriched in aging in the brain, heart, muscle, and spleen. The top 5 GO terms that are enriched in aged tissue (top, red) and the top 5 terms that are enriched in young tissue (bottom, off-white) are shown and ordered by significance. All tissues have at least one term associated with immunity enriched with age (bold). (B) Top GO terms enriched in each sex in the brain, heart, muscle, and spleen. The top 5 GO terms enriched in females (top, pink) and top 5 GO terms in males (bottom, blue) are shown and ordered by significance. The spleen had the least female-biased sexual dimorphic gene expression with only one term significantly enriched (FDR < 5%).

### Usage Notes

In this study, we generated ribosomal RNA-depleted RNA-seq data of 4 tissues from young and old turquoise killifish from the GRZ strain across both sexes. Importantly, this is the first turquoise killifish RNA-seq dataset to profile both aging and sex across multiple tissues using the reference (GRZ) strain. Turquoise killifish RNA-seq studies to date have focused on either sex or aging independently, generally in a single tissue, or conducted the study in a non-reference strain of turquoise killifish^2–8^.

In our dataset, we observed in all tissues analyzed underwent greater transcriptional changes with aging, than between males and females. This pattern held true for both genes and TEs. TE expression was very high in the brain, with roughly half of all reads in each library mapping to TE sequences. Why the brain is so amenable to TE expression remains a mystery, though a role of TEs in memory formation may play a part^46^. Furthermore, the brain had a dramatic upregulation of TEs with age. Similar findings have been reported in other animals including Drosophila and mice and may be reflective of a conserved mechanism of brain aging^7, 47–49^.

The results of the GO analysis suggest that killifish tissues experience an increase in immune-related transcription with age. This phenomenon, referred to as “inflamm-aging” is a well described aspect of aging and helps validate our dataset^45^. Furthermore, we find that terms related to cell division decrease in expression in aged muscles. This may capture a decrease in muscle stem cell numbers, which is reported to occur in the turquoise killifish^50^. Sex-dimorphic GO terms appear to be less conserved across tissues and may be a rich area for further exploration of the differential effects of sex on transcription.

This dataset can be used to find differences in gene and TE expression using age and sex as variables in any combination suitable to the user. In addition, to facilitate the exploration of this dataset, we have deployed a user-friendly searchable database of differential gene and TE expression results, with human homology information, that can be mined by the community (https://alanxu-usc.shinyapps.io/nf_interactive_db/). The dataset could also be deconvoluted using single-cell atlases to establish cell composition profiles and analyze how cell type frequencies change with age and sex in each tissue.

Since the dataset was generated using ribosomal RNA depletion rather than polyA enrichment, it should also be possible to analyze RNA species other than canonical mRNAs, including circRNAs^51^ transcribed by RNA pol III, which typically lack polyadenylation^52, 53^.

Limitations of this dataset are that, like most aging −omic studies outside of consortia efforts, it uses only 2 timepoints, which limits ability to enable detection of specific changes at middle-age^17^. In addition, TE quantification may be partially driven by TE-derived intronic reads that are retained by ribosomal RNA-depleted RNA-seq library preparation^54^. In effect, this dataset cannot distinguish between intronic-derived TEs and autonomous TEs, which are regulated in a different fashion, although both may contribute to biological changes. Nonetheless, this dataset is useful in determining the total amount and class of TE reads present in young and old tissues across sexes.

### Code Availability

All analytical code used for processing and technical validation is available on the Benayoun Laboratory GitHub repository (https://github.com/BenayounLaboratory/Killifish_RNASeq_2023). The provided R code was run and tested on R v4.3.0.

## Acknowledgments

Some figure elements were generated with BioRender (https://biorender.com) and Freepik. We would like to acknowledge the Center for Advanced Research Computing (CARC) at USC for providing the computational resources used to perform many of the analyses used in this study.

This work was supported by a National Institute on Aging (NIA) T32 AG052374 postdoctoral training grant fellowship to B.B.T., grant R35 GM142395 from National Institute of General Medical Sciences, a pilot grant from the NAVIGAGE Foundation, and a Hanson-Thorell Family award to B.A.B.

## Author Contributions

Conceptualization A.X., D.R.V and B.A.B; animal husbandry and dissection D.R.V., A.T.; RNA isolation S.N.; RNA-seq library preparation R.J.L., S.N.; data analysis A.X., B.B.T, B.A.B; manuscript preparation A.X, B.B.T., B.A.B. Manuscript editing: all authors.

## Completing Interests

The authors declare no competing interests.

## Figures

Figures 1-6 provided as a separate file (pdf).

